# Unraveling tumor heterogeneity: Quantitative insights from scRNA-seq analysis in breast cancer subtypes

**DOI:** 10.1101/2024.08.30.610531

**Authors:** Daniela Senra, Nara Guisoni, Luis Diambra

**Affiliations:** Centro Regional de Estudios Genómicos, Universidad Nacional de La Plata, Argentina; Departamento de Física, Facultad de Ciencias Exactas, Universidad Nacional de La Plata

**Keywords:** Breast cancer, scRNA-seq, Tumor heterogeneity, Copy number alteration

## Abstract

Tumors are complex systems characterized by genetic, transcriptomic, phenotypic, and microenvironmental variations. The complexity of this heterogeneity plays a crucial role in metastasis, tumor progression, and recurrence. In this work, we utilized publicly available single-cell transcriptomics data from human breast cancer samples (ER+, HER2+, and triple-negative) to evaluate key concepts pertinent to cancer biology. Quantitative assessments included measures based on copy number alterations (CNAs), entropy, transcriptomic heterogeneity, and different protein-protein interaction networks (PPINs).

We found that entropy and PPIN activity related to the cell cycle delineate cell clusters with notably elevated mitotic activity, particularly elevated in aggressive breast cancer subtypes. Additionally, CNA distributions differentiate between ER+ and HER2+/TN subtypes. Further, we identified positive correlations among the CNA score, entropy, and the activities of PPINs associated with the cell cycle, as well as basal and mesenchymal cell lines. These scores reveal associations with tumor characteristics, reflecting the known malignancy spectrum across breast cancer subtypes.

By bridging the gap between existing literature and a comprehensive quantitative approach, we present a novel framework for quantifying cancer traits from scRNA-seq data by establishing several scores. This approach highlights the potential for deeper insights into tumor biology compared to conventional marker-based approaches.

## 1. Introduction

Breast cancer is the most commonly diagnosed cancer and the fifth leading cause of cancer mortality worldwide [1,2]. It comprises a diverse set of diseases, leading to the utilization of various classification systems over time. Currently, immunohistochemistry of hormone receptors, including the estrogen receptor (ER), progesterone receptor (PR), and human epidermal growth factor receptor 2 (HER2), is one of the most extensively employed approaches for classification. This system identifies four main subtypes of breast cancer: luminal A, luminal B, HER2-amplified, and triple-negative [3–5].

Luminal A and B subtypes, defined by the presence of estrogen receptors (ER+), exhibit distinct characteristics. Luminal B shows reduced ER and PR levels, higher proliferation rates, and variable HER2 expression compared to Luminal A, which typically expresses low HER2 levels [6,7]. Together they constitute approximately 70% of all breast cancers and generally carry a favorable prognosis. HER2-amplified (HER2+) breast cancer is characterized by heightened HER2 expression and absence of ER. HER2+ breast cancer represents between 15 − 20% of breast cancers and manifests a more aggressive behavior compared to the luminal subtypes [4]. Triple-negative (TN) breast cancer makes up to 15% of breast cancer diagnoses, and lacks ER, PR, and HER2 expression, demonstrating the most aggressive behavior with poor differentiation and elevated proliferation rates. The terms TN and basal breast cancer are frequently used interchangeably due to significant similarities in their signatures; however, not all TN breast cancers are basal [8,9].

The advent of single-cell RNA sequencing (scRNA-seq), enabling transcriptomic profiling of individual cells, has introduced new perspectives. scRNA-seq offers the advantage of unraveling the complex composition of mixed populations within a sample compared to traditional bulk analysis. Several human breast cancer datasets, spanning from a few hundred to hundreds of thousands of cells, have become available [10–13]. Significant efforts have been dedicated to dissecting cell heterogeneity and identifying cell types and states through clustering analysis [14–16].

In this study, we adopt a quantitative methodology to comprehensively investigate several classical cancer-associated hallmarks from breast cancer scRNA-seq data. These include intratumor transcriptomic heterogeneity, copy number alterations (CNAs), entropy, and the activity of protein-protein interaction networks (PPINs) of interest. We evaluate these features by establishing mathematical scores. While previous research has examined these features separately across various cancer types, such investigations often relied on qualitative or semi-quantitative assessments. Below, we outline the features studied in this work and provide a brief overview of the current state of research in each area.

### Intratumor transcriptomic heterogeneity

Tumors exhibit a wide range of phenotypic and molecular characteristics, at both intertumor and intratumor levels. Heterogeneity, extensively studied for its implications in diagnostics and treatment selection, poses significant challenges, as different tumors and cells respond differently to the same therapeutic approach [17]. Intertumor heterogeneity encompasses variations observed between distinct tumors from different or the same patient. Furthermore, a single tumor comprises a mixture of tumor cell subpopulations with diverse genetic, transcriptomic, and phenotypic profiles. This phenomenon, known as intratumor heterogeneity, is the consequence of clonal evolution and the influence of diverse microenvironmental factors[17,18]. Intratumor heterogeneity presents as a selective benefit, increasing the probability of both the pre-existence of tolerant and resistant subpopulations and the ability to adapt [19].

In recent years, several studies have assessed heterogeneity at the transcriptomic level in breast cancer using scRNA-seq data [10–13,20]. These investigations have focused on delineating heterogeneity through the expression profiling of markers associated with breast cancer, as well as identifying distinct cell clusters and their corresponding gene expression signatures. However, while quantitative assessments of intratumor heterogeneity have been conducted in various other cancer types [11,21], the field of breast cancer research lacks a quantitative exploration of intratumor heterogeneity.

### CNA

CNAs are changes in the number of gene copies in tumor cells and are thought to give a selective advantage to the cells within tumors, resulting in increased expression of certain genes and reduced expression of others. Tumors utilize DNA loss as a mechanism to eliminate tumor suppressor gene copies, whereas gains in DNA copy numbers can increase the activity of oncogenes, promoting tumor progression. CNAs are a distinguishing feature of human cancers, reflecting the extent and type of genomic instability unique to each [22]. It has been reported that CNAs are linked with cancer progression and poor prognosis in breast cancer [23,24]. In previous works, some scores were defined to account for the extent of CNAs at the cell level from scRNA-seq data in pancreatic cancer and glioblastoma [25,26]. However, in the context of breast cancer, CNA inference has primarily been focused on discriminating cancer cells from non-tumor cells, rather than quantifying the extent of CNAs [10,11].

### Entropy

Entropy is a quantity that describes the degree of disorder or uncertainty in physical systems. This concept has been adapted to different disciplines including the field of single-cell transcriptomics. Entropy and entropy-based metrics have served as valuable tools for quantifying cell heterogeneity and other attributes, including the assessment of stemness [27,28]. In the single-cell transcriptomics field, entropy reflects the diversity or variability in gene expression across individual cells. Highly specialized cells typically express a specific set of genes associated with their biological function, leading to lower entropy levels. In contrast, cells with less defined roles tend to display a wider range of active genes simultaneously, resulting in higher entropy levels. Banerji *et al*. computed an entropy-based metric from bulk data on breast cancer and observed that patients with a worse prognosis exhibited higher scores. This metric proved to be a more robust predictor of prognosis compared to other commonly used prognostic indicators [29].

### PPIN activity

Enrichment of gene sets allows to explore the expression of relevant pathways of in scRNA-seq data [30,31]. Cell cycle and proliferation are associated with the malignancy of breast cancer and, marker genes associated with those processes, such as MKI67, are usually used as indicators of prognosis [32,33]. Breast cancer scRNA-seq datasets have revealed clusters of highly proliferating cells, and the proportion of these cells correlates with cancer aggressiveness [10,11]. In breast cancer, epithelial to mesenchymal transition (EMT) has been reported to be associated with cancer progression [34]. However, unlike proliferation, Pal *et al*. did not observe a cluster of cells with increased expression of EMT markers in breast cancer scRNA-seq datasets [10]. In another study using single-cell spatial transcriptomics, a group of spatially contiguous cells demonstrated enrichment of EMT markers [11]. Much remains unknown about the EMT program in breast cancer and better markers need to be explored [35]. Furthermore, the inherent sparsity of scRNA-seq data poses challenges when attempting to quantify the levels of biological processes using markers. An alternative to address this limitation is the usage of PPINs associated with a set of genes, such as those involved in specific biological processes [36]. In this manner, regulatory information is also taken into consideration, providing measurements of the activity levels of PPINs that are less sensitive to the zero-inflated expression matrices. Consequently, pathways of interest, such as EMT or cell cycle, can be assessed from scRNA-seq data.

This work presents a novel approach to the quantitative analysis of these fundamental concepts, specifically applied to a breast cancer scRNA-seq dataset. By quantifying these features and investigating their associations with the three primary breast cancer subtypes (ER+, HER2+, and TN), our research aims to deepen the understanding of the intricate relationships between these features and breast cancer subtypes.

## 2. Materials and methods

### 2.1 Dataset description

The dataset utilized in this study was obtained from scBrAtlas [10], which is available in two forms: raw count matrices in the GEO database and preprocessed R objects on figshare [37]. For our analysis, we employed the preprocessed R objects from figshare.

The scBrAtlas is a collection of scRNA-seq samples derived from various human breast states, including normal, preneoplastic and cancerous conditions. This atlas encompasses around 430000 individual cells, obtained from a total of 69 surgical samples collected from 55 patients.

The normal breast samples were procured from reduction mammoplasties performed on donors with no family history of breast cancer. The preneoplastic samples, on the other hand, were obtained from individuals carrying the BRCA1 mutation, a genetic predisposition associated with an increased risk of developing breast cancer. The cancer samples were sourced from primary tumors of treatment-naïve patients encompassing three major subtypes: ER+, HER2+ and TN. These samples contained epithelial cells (normal and cancer), stromal, and immune microenvironment cells.

### 2.2 Analysis of scRNA-seq data

Quality control metrics were obtained to ensure the reliability of the data. The samples underwent a filtering process based on criteria such as library size, number of genes per cell, and the percentage of mitochondrial content per cell. As a result of this trimming process, approximately 15% of the cells were filtered out for each sample, leaving a total of 341874 cells for further analysis. A detailed description of the preprocessing phase, along with the corresponding R code, can be found in [38]. Since the data had already undergone preprocessing, this study verified this step and used directly the preprocessed data as is available.

Our primary interest was the study of cancer cells, therefore we exclusively utilized samples obtained from women with breast cancer. Male, healthy, precancerous, and cancer samples associated with lymph nodes were not considered for further analysis. The age of the donors ranges from 25 to 84 years old. In Table 2.1 a detailed description of the samples is provided, including patient age, cancer subtype, size, grade, and number of cancer cells. All samples consisted of a mixture of tumor cells, normal epithelial cells, and cells from the microenvironment such as fibroblasts, endothelial cells, and immune cells, among others. For downstream analysis, we excluded all cells from the microenvironment, retaining only epithelial cells. Furthermore, we subsetted the cancer cells, filtering out normal epithelial cells. The cells were initially labeled by the original data source, with authors distinguishing cancer from normal cells using inferCNV [39–41]. To validate the provided cell labels and confirm the distinction between tumor and normal cells, we independently ran inferCNV. After performing cell filtering, we excluded samples containing fewer than 1000 cancer cells.

**Table 1.** Description of breast cancer samples. Samples from male patients and those associated with lymph nodes have been omitted from the listing, as they were not included in the analysis. The samples selected for analysis, meeting the criterion of having more than 1000 tumor cells after filtering, are marked with an asterisk next to their respective sample IDs. The table includes details on Sample ID, patient age, cancer subtype, size, grade, and the number of cancer cells that remained after applying filtering. The complete metadata can be found in the original publication associated with the data in the *Supporting Information* section as Table EV2, Table EV3 and Table EV4 [10].

| Sample ID | Patient age | Cancer subtype | size (mm) | Grade | cancer cells after filtering |
| --- | --- | --- | --- | --- | --- |
| TN-0126* | 64 | TN | 64 | 3 | 1235 |
| TN-0135* | 61 | TN | 22 | 3 | 14333 |
| TN-0106 | 65 | TN | 25 | 3 | 54 |
| TN-0114-T2 | 84 | TN | 17 | 3 | 177 |
| TN-B1-4031* | 25 | TN (BRCA1) | 20 | 3 | 5129 |
| TN-B1-0131* | 84 | TN (BRCA1) | 25 | 3 | 5513 |
| TN-B1-0554* | 29 | TN (BRCA1) | 37 | 3 | 2337 |
| TN-B1-0177* | 30 | TN (BRCA1) | 13 | 3 | 1575 |
| HER2-0308* | 32 | HER2+ | 20 | 3 | 3317 |
| HER2-0337* | 66 | HER2+ | 67 | 3 | 3924 |
| HER2-0031* | 47 | HER2+ | 18 | 3 | 1606 |
| HER2-0069 | 71 | HER2+ | 27 | 3 | 196 |
| HER2-0161* | 80 | HER2+ | 45 | 3 | 4124 |
| HER2-0176* | 60 | HER2+ | 20 | 3 | 4682 |
| ER-0319 | 58 | PR+ | 27 | 3 | 568 |
| ER-0001* | 58 | ER+ | 32 | 3 | 4559 |
| ER-0125* | 45 | ER+ | 48 | 2 | 3678 |
| ER-0360* | 70 | ER+ | 50 | 2 | 1934 |
| ER-0032 | 55 | ER+ | 90 | 3 | 417 |
| ER-0042* | 58 | ER+ | 18 | 2 | 2899 |
| ER-0025* | 52 | ER+ | 23 | 2 | 4499 |
| ER-0151 | 49 | ER+ | 35 | 2 | 749 |
| ER-0163* | 45 | ER+ | 45 | 3 | 5378 |

**Table 2.**
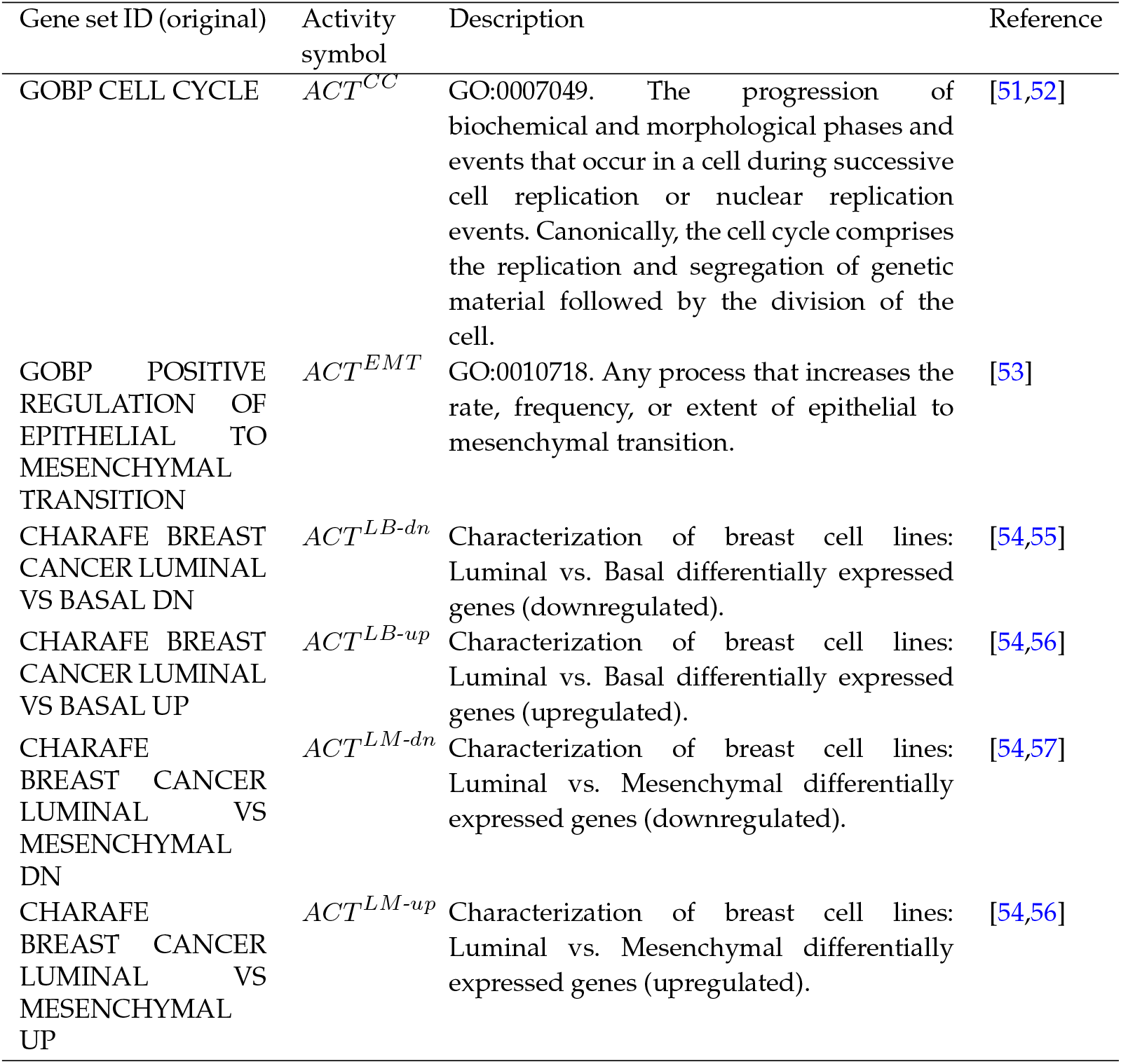
Details regarding the gene sets for which the protein-protein interaction network (PPIN) activity was calculated.

The complete dataset [10] incorporated a total of nineteen ER+, six HER2+, and eight TN breast cancer samples. As per these selection criteria, six ER+ samples (ER-0001, ER-0125, ER-0360, ER-0042, ER-0025 and ER-0163), five HER2+ samples (HER2-308, HER2-0337, HER2-0161 and HER2-0176) and six TN samples (TN-0126, TN-0135, TN-B1-4031, TN-B1-0131, TN-B1-0554 and TN-B1-0177) were considered for subsequent downstream analysis. The names of the samples provided by the data authors were maintained.

We used R (version 4.3.1) and the Seurat package (version 4.4.0) for data preprocessing and data analysis. Samples were integrated separately across sample subtypes using the Seurat 4 pipeline [42,43]. In figures 1A-C, ER+, HER2+, and TN breast integrated datasets are shown and samples are distinguished by the color of the cells.

**Figure 1.**
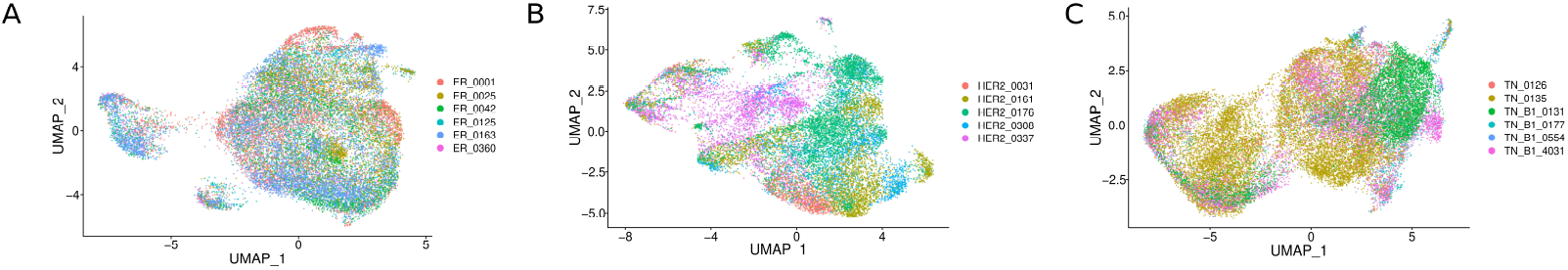
UMAP visualization of integrated samples separated by cancer subtype, ER+ (A), HER2+ (B), and TN (C). Cells are color-coded according to the sample.

### 2.3 Scores

To investigate potential links between tumor aggressiveness and specific biological features, we established several parameters to measure the extent of these features. These parameters include the level of CNAs, intratumor heterogeneity, entropy, and the activity of specific PPINs potentially associated with tumor aggressiveness, such as EMT and cell cycling. In this section, we provide precise definitions of these measures and the rationale for their application. All the relevant codes to calculate the scores are freely available at https://github.com/danielasenraoka/R-code-workflow-Unraveling-tumor-heterogeneity and the code to calculate activity of PPIN’s is available at https://github.com/danielasenraoka/PyOrigins.

#### 2.3.1 CNA score

One of our main interests was looking at somatic CNAs. To deduce CNAs from scRNA-seq data, we employed the inferCNV R package (version 1.16.0) developed by the Trinity CTAT Project [39–41]. InferCNV serves as a tool for the detection of somatic large-scale chromosomal CNAs, including entire chromosomal gains or deletions and substantial segments of chromosomes. This is achieved by assessing the relative gene expression levels of contiguous genes along the genome and comparing them to a reference set of “normal” cells. This approach is a well-established method utilized for distinguishing between tumor and normal cells.

In our study, inferCNV was applied separately to samples based on cancer subtypes. We used a breast normal epithelial sample labeled as N0372, as our reference cell population, as was also done by the dataset creators [10]. A sliding window of 100 contiguous genes was employed. The pre-existing classification of cells into cancer and non-cancer categories, as provided alongside the preprocessed dataset, was validated using the inferCNV profiles obtained in alignment with the labeled data.

Computational tools, including inferCNV, copyKAT [44], and CaSpER [45] are commonly used to estimate CNAs in scRNA-seq data. These tools are primarily designed to distinguish tumor cells from non-tumor cells. However, they can also be valuable for quantifying the extent of CNA within individual cells. To achieve this, we define a CNA level for each cell *i* based on the residual expression matrix generated by inferCNV:

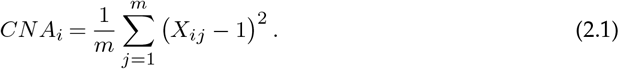

We denote the transposed residual expression matrix obtained from inferCNV by *X*, a matrix with *n* rows (cells) and *m* columns (genes). We avoid using the transposed symbol to simplify notation. This matrix serves as a surrogate indicator for CNA in each gene *j* across every cell *i*. A value of 1 signifies neutrality, values exceeding 1 indicate gains, and values below 1 denote losses. Hence, all elements of the matrix are standardized by subtracting 1. It is important to highlight that since the terms are squared, the score reflects the magnitude without distinguishing between gains and losses. This score indicates the mean squared dispersion of CNAs relative to the reference normal cells, and it is a variation of scores defined in prior works [25,26].

To quantify the degree of sample CNA burden the *CNA* score can be averaged across all the cells within the sample as follows:

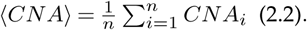

#### 2.3.2 Transcriptomic heterogeneity

The standard approach for evaluating intratumor heterogeneity in cancer using scRNA-seq data involves performing clustering to identify distinct cell types or states [46,47]. Subsequently, these clusters are characterized using differential expression analysis and enrichment analysis to obtain gene markers for each cluster. While this method yields valuable insights into sample composition, it does not provide a score that measures the extent of heterogeneity at a transcriptomic level. To address this, we propose an alternative to assess transcriptomic heterogeneity inspired by previous works [43].

Our proposed approach involves assessing the variability of each gene from a scRNA-seq sample. Deriving variance from the log-normalized data fails to account for the inherent mean-variance relationship present in scRNA-seq data. To address this mean-variance relationship, variance-stabilizing methods are typically used [48,49]. In our study, we utilized the variance-stabilizing method from the Seurat package, specifically employing the FindVariableFeatures and HVFinfo functions, which provide an estimator of variance adjusted for the feature mean [42]. This approach allows us to accurately assess gene variability while accounting for the underlying mean-variance relationship in scRNA-seq data. To quantify this variability across the entire transcriptome, we define a sample expression variability score as:

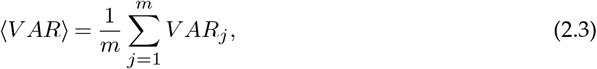

where *V AR*_*j*_ represents the variance stabilized of gene *j*. ⟨*V AR*⟩ denotes the mean-variance across genes in a sample, serving as a metric to assess the degree of intratumor heterogeneity within a sample.

This metric quantifies intratumor heterogeneity by calculating the average standardized variance across all expressed genes. A lower score indicates greater transcriptomic homogeneity, meaning cells within a sample share more similar gene expression profiles. Conversely, higher values signify increased transcriptomic heterogeneity, suggesting greater diversity among the cells.

#### 2.3.3 Activity of protein-protein interaction networks

In previous works, we have developed a methodology to quantify the cell activity of a PPIN from scRNA-seq data [36,50]. A PPIN associated with a specific set of genes can be constructed by extracting the interactions from the full Human PPIN that involve proteins encoded by the genes within the defined gene set *x*. A score that reflects the activity of a PPIN can be defined for each cell *i* as follows:

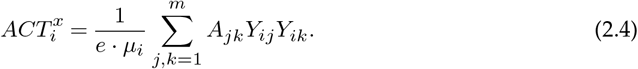

Where *A*_*jk*_ is the upper triangular adjacency matrix, that characterizes the connectivity between genes *j* and *k* in the network. *Y* is the transposed normalized expression matrix of dimensions *n* × *m, e* is the number of edges in the PPIN, and *μ*_*i*_ is the average expression of all genes in cell *i*. To enhance clarity, each row of *Y* is the expression profile of the *i*-th cell. Hence, *Y* is the transposed counterpart of the conventional expression matrix, where rows correspond to genes and columns to cells. Again, this adjustment enhances clarity regarding notation to avoid using transposed symbols or non-standard index orders. A normalization factor of *e* · *μ*_*i*_ is introduced to account for graph size differences and differences in the mean expression. This factor enables comparisons between samples and various PPINs.

To quantify the overall activity levels of a PPIN within a sample, we can compute the average activity across all cells:

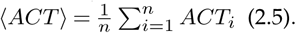

We computed the activity of the PPINs associated with six Homo sapiens gene sets as detailed in Table 2. Genes associated with the biological process cell cycle (GO:0007049) were sourced from the QuickGO database [51,52] and negative regulators of this process were excluded as was previously done [36]. The remaining gene sets were retrieved from the Molecular Signatures Database (MSigDB) [58,59]. These include positive regulation of EMT and four gene sets associated with a study that characterizes different breast cancer cell lines (basal, luminal, and mesenchymal) [54]. These gene sets provide comprehensive insights into distinct cellular behaviors across different breast cancer subtypes. The activities of the PPINs associated with the cell cycle and EMT are denoted as *ACT*^*CC*^ and *ACT*^*EMT*^ respectively. We refer to *ACT*^*LB*-*up*^, *ACT*^*LM*-*up*^, *ACT*^*LB*-*dn*^, and *ACT*^*LM*-*dn*^ as the PPIN activities corresponding to the up-regulated and down-regulated differentially expressed genes between luminal vs. basal and luminal vs. mesenchymal breast cancer cell lines.

#### 2.3.4 Entropy

We compute the Shannon entropy, a well-established information theory metric used to measure the degree of miss-information of a system configuration. Suppose we are working with a scRNA-seq dataset, represented by the transposed count matrix denoted as *Z*. In this context, we can calculate the Shannon entropy of a specific cell *i* as follows:

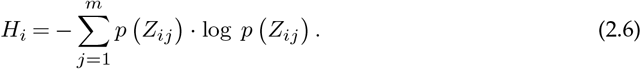

Entropy is associated with a probability mass function *p*(*Z*_*ij*_) of cell *i* expressing gene *j*. In scRNA-seq data, it can be estimated by dividing the expression of gene *j* by the total expression of that particular cell: 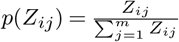 [27,60,61]. To quantify the degree of entropy for a sample, we can simply average entropy across all cells comprising the sample as follows:

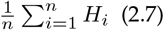

Traditionally linked to heterogeneity, in this context entropy measures the heterogeneity of individual cells, not the sample intratumor heterogeneity. High *H* indicates a broader range of genes expressed simultaneously within a single cell, a characteristic of non-specialized cells such as stem or progenitor cells [27,62]. In our study, the score ⟨*H*⟩ represents the average entropy across all cells within a sample. Hence, samples with higher ⟨*H*⟩ scores exhibit more undifferentiated characteristics associated with cancer stemness. It has been reported that breast cancer aggressiveness and therapy resistance may be driven by breast cancer stem cells (CSCs) [63–65]. Notably, it was observed that in TN tumors CSCs are enriched compared to non-TN breast cancers [66].

In summary, we established parameters derived from scRNA-seq data, which capture the levels of CNAs, entropy, and the activity of PPINs at the individual cell level. Additionally, we assessed the transcriptomic intratumor heterogeneity at the sample level. Overall cell score parameters at the sample level were derived by averaging across all cells within each sample.

## 3. Results and discussion

We examined 17 individual samples obtained from breast cancer patients (refer to Table 2.1). These samples were categorized by cancer subtype and integrated (figures 1A-C). To delve into CNA, we conducted separate CNA analyses for each cancer subtype. The resulting heatmaps of chromosomal copy gains or losses are depicted in figures 2A-C.

**Figure 2.**
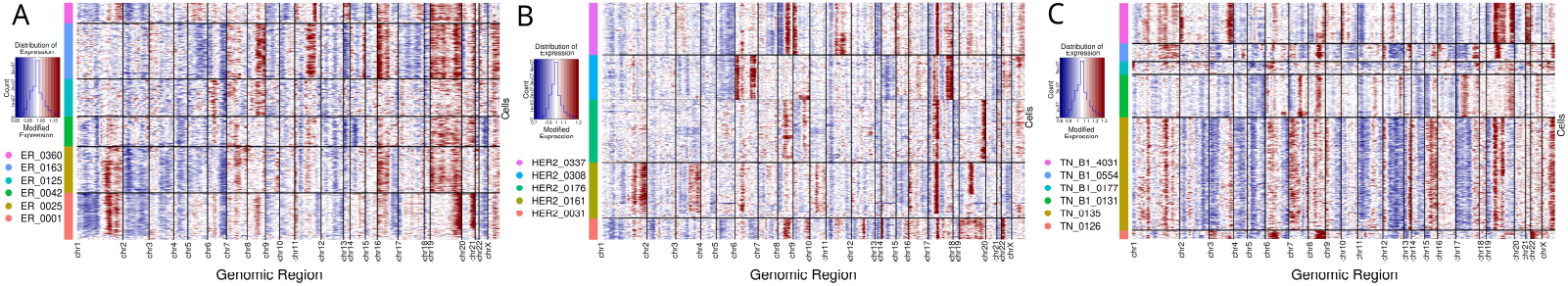
InferCNV heatmap plots displaying inferred CNAs for each cancer subtype, ER+ (A), HER2+ (B), and TN (C). Columns depict genes sorted by genome location and grouped by chromosomes, while rows represent individual cells grouped by samples and color-coded according to the respective sample. Amplifications are indicated in red and deletions in blue.

Although the gain-deletion patterns do not present great synteny between the subtypes, some common characteristics can be highlighted. Chr1 exhibited gains at chr1q, which contains several oncogenes such as NRAS, JUN, MYCL, TAL1, and BLYM, and losses at chr1p. Notably, chr1q gains have been reported in approximately 60% of patients, primarily associated with ER+ cancer [67]. Deletions in chr2 were observed in nearly all samples, irrespective of subtype. Chr8q amplifications, encompassing the MYC proto-oncogene, were frequent across all subtypes, confirming their well-documented role in breast cancer [68–70]. Chr19 amplifications were seen in most samples, but were more pronounced in ER+ and TN subtypes, aligning with previous findings [71]. Across each subtype, common characteristics among patients are noted. HER2+ samples displayed consistent chr17 amplifications, particularly at the chr17q12 band harboring the HER2 gene, and frequent chr13 deletions. Despite identifying these recurrent *CNAs* across subtypes, significant heterogeneity within each subtype was observed. This phenomenon, previously reported in breast tumors [68], underscores the complex and diverse nature of these malignancies. Note that samples HER2-0308 and TN-B1-0131 exhibit CNA profiles that differ from their respective subtype patterns. This aspect will be detailed discussed ahead.

To delve into the heterogeneity of tumors, we analyzed each cancer subtype using the scores defined in the Methods section. The cell distributions of eight derived scores for each sample can be seen in violin plots of figure 3. Additionally, figure 4 visualizes the distribution of a subset of scores for individual cells within the integrated UMAP space, specific to each of the three cancer subtypes.

**Figure 3.**
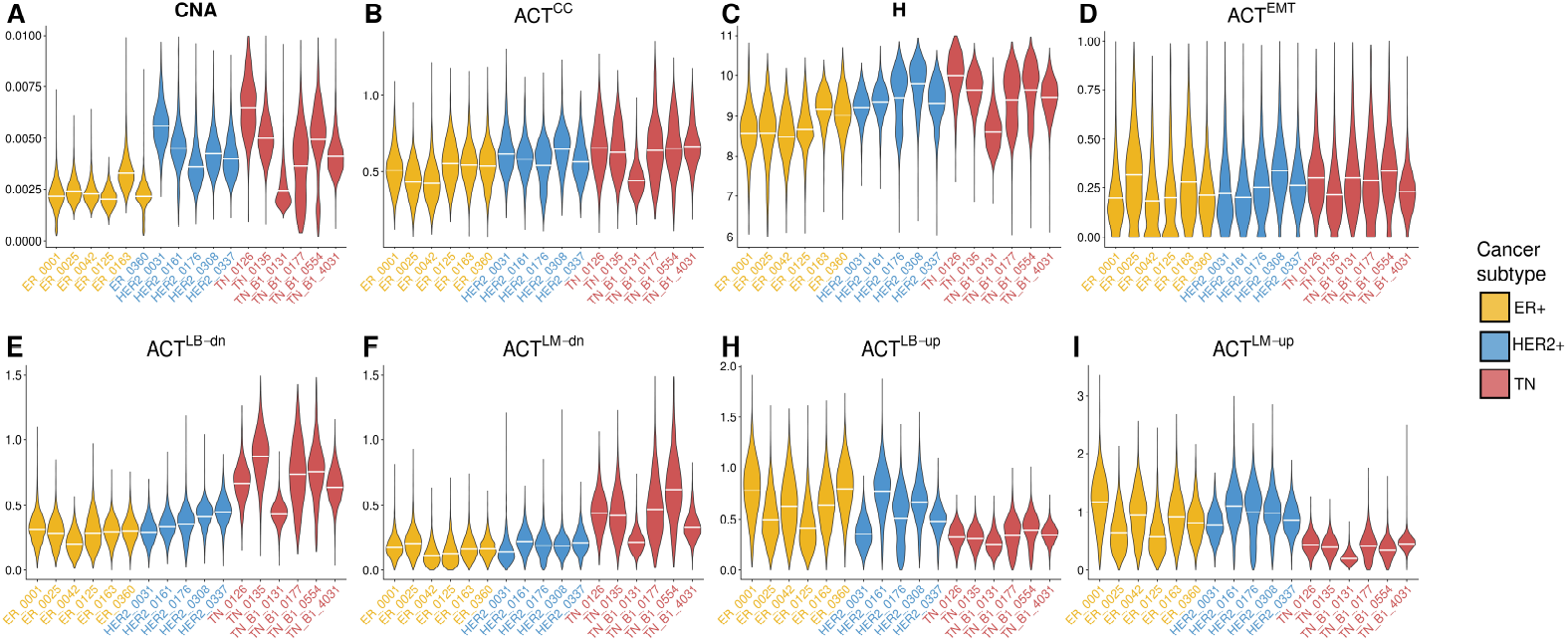
(A-I) Cell distribution of *CNA, ACT*^*CC*^, *H, ACT*^*EMT*^, *ACT*^*LB*-*dn*^, *ACT*^*LM*-*dn*^, *ACT*^*LB*-*up*^, and *ACT*^*LM*-*up*^ per sample. The white horizontal lines represent the mean value of the corresponding score for each sample. Yellow, blue, and red sample labels correspond to ER+, HER2+, and TN samples respectively.

**Figure 4.**
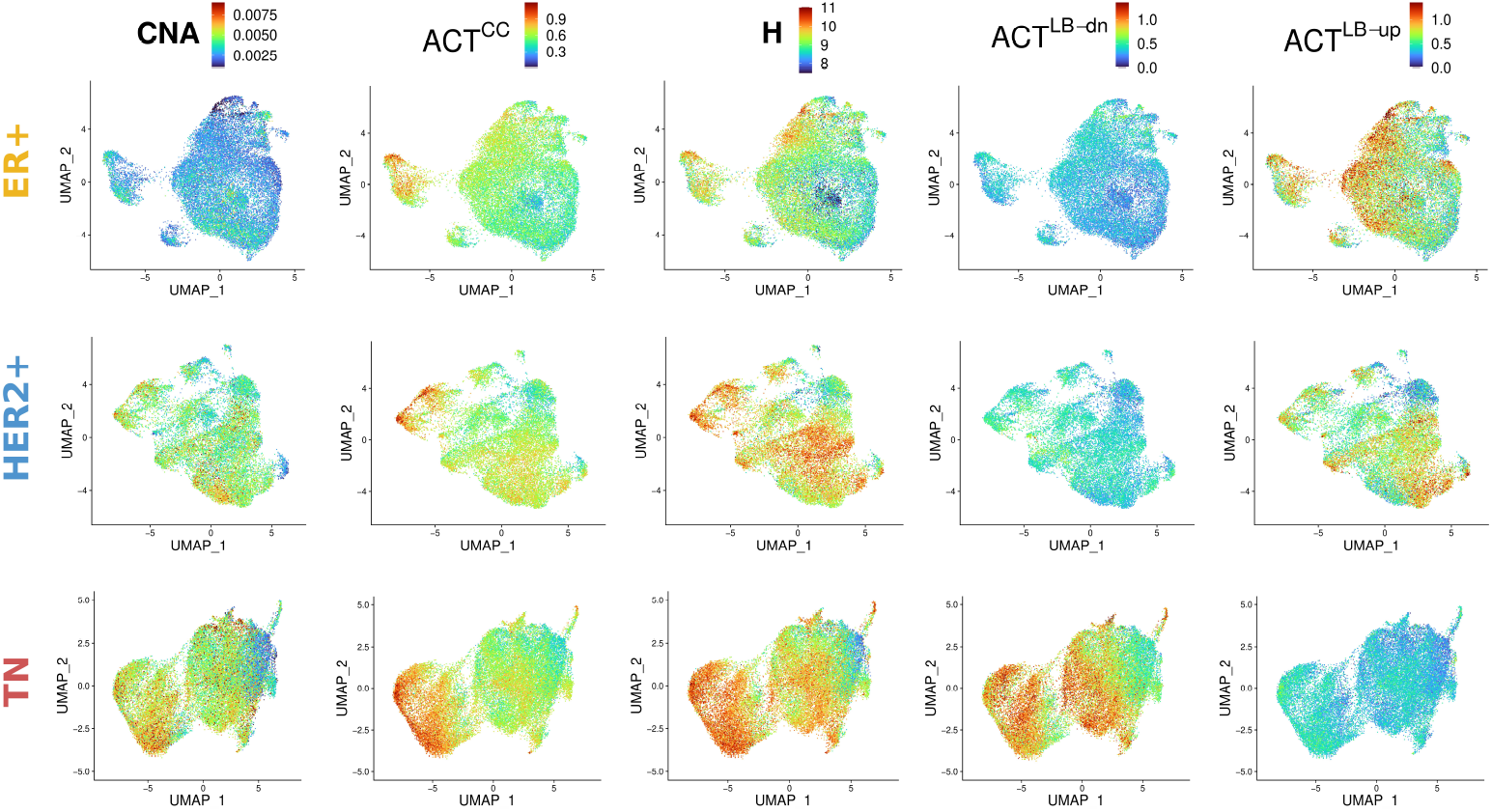
UMAP representation of integrated samples separated by cancer subtype: ER+, HER2+, and TN. Cells are color-coded according to the scores *CNA, ACT*^*CC*^, *H, ACT*^*LB*-*dn*^, and *ACT*^*LB*-*up*^.

The distributions of the *CNA* score are illustrated in figure 3A, revealing lower values in ER+ samples compared to those observed in HER2+ and TN samples (Mann-Whitney Test, *p*-values= 0.00466 and 0.00824, respectively). Furthermore, no distinct cell clusters based on this score are observed within the UMAP embedding (see figure4). In contrast, as can be seen in figures 3B and C, the scores ⟨*ACT*^*CC*^⟩ and ⟨*H*⟩ show a gradual increase across ER+, HER2+, and TN samples. In line with previous works [10,38], cell clusters exhibiting high *ACT*^*CC*^ were observed across all cancer subtypes, as depicted in figure 4. The discrete clusters of cycling MKI67+ tumor cells identified by the dataset authors coincided with cell clusters exhibiting elevated *ACT*^*CC*^ across all cancer subtypes. Furthermore, consistent with these studies, TN samples demonstrated a higher proportion of cells with elevated *ACT*^*CC*^ compared to ER+ and HER2+ cancer subtypes.

The distribution of *ACT*^*EMT*^ across samples, depicted in figure3D, shows no significant differences (Mann-Whitney Test, *p*-values > 0.065). Furthermore, this score does not present evidence of cell clusters exhibiting higher levels of activity in this PPIN (see sup figure S1), in agreement with previous observations [10].

Additionally, we explored the activity of the PPINs associated with breast cancer cell lines [54] (see Method Section). *ACT*^*LM*-*up*^ and *ACT*^*LM*-*dn*^ distributions in the UMAP space were similar to *ACT*^*LB*-*up*^ and *ACT*^*LB*-*dn*^, respectively (see figures 4 and S1). As expected, TN samples display higher values of *ACT*^*LB*-*dn*^ and *ACT*^*LM*-*dn*^ scores than ER+ samples (Mann-Whitney Test, *p*-value=0.00507 in both cases), as illustrated in figures 3E, F, and 4. Conversely, as can be seen in figures 3H, I, and 4, ER+ samples exhibit significantly higher *ACT*^*LB*-*up*^ and *ACT*^*LM*-*up*^ scores compared to samples derived from basal/mesenchymal tumors, i.e. TN samples (Mann-Whitney Test, *p*-value = 0.00305 in both cases). In a similar manner the samples derived from the other luminal tumor (HER2+) also exhibit significantly higher *ACT*^*LB*-*up*^ and *ACT*^*LM*-*up*^ scores compared to TN samples, (Mann-Whitney Test, *p*-values = 0.00811 and 0.00466, respectively). These results indicate that many tumor cells preserve the transcriptional landscape of the original lineage. Furthermore, ER+ and HER2+ samples display cell clusters with high *ACT*^*LB*-*up*^ scores, whereas TN samples exhibit clusters with high *ACT*^*LB*-*dn*^ scores (figure 4). Interestingly, cells with strong original lineage features are co-localized in the UMAP space with cells associated with high entropy *H*.

To assess the relationship between the scores we computed the correlation matrix among sample averages using the Pearson correlation coefficient, depicted in figure 5A. At the sample level, the mean scores ⟨*CNA*⟩, ⟨*ACT*^*CC*^⟩, ⟨*H*⟩, ⟨*ACT*^*LB*-*dn*^⟩, ⟨*ACT*^*LM*-*dn*^⟩, ⟨*ACT*^*EMT*^⟩, and ⟨*V AR*⟩ revealed positive correlations. Notably, the first five scores exhibited stronger correlations among themselves forming a cluster obtained by performing hierarchical cluster analysis (hclust from stats R package version 4.3.1), as highlighted in figure 5A. Specifically, ⟨*CNA*⟩, ⟨*H*⟩, and ⟨*ACT*^*CC*^⟩ displayed the strongest Pearson correlation coefficients, ranging from 0.77 to 0.88. Moreover, both ⟨*V AR*⟩ and ⟨*ACT*^*EMT*^⟩ exhibited positive correlations with these scores. Conversely, ⟨*ACT*^*LM*-*up*^⟩ and ⟨*ACT*^*LB*-*up*^⟩ demonstrated negative correlations with all other scores, except for ⟨*V AR*⟩.

**Figure 5.**
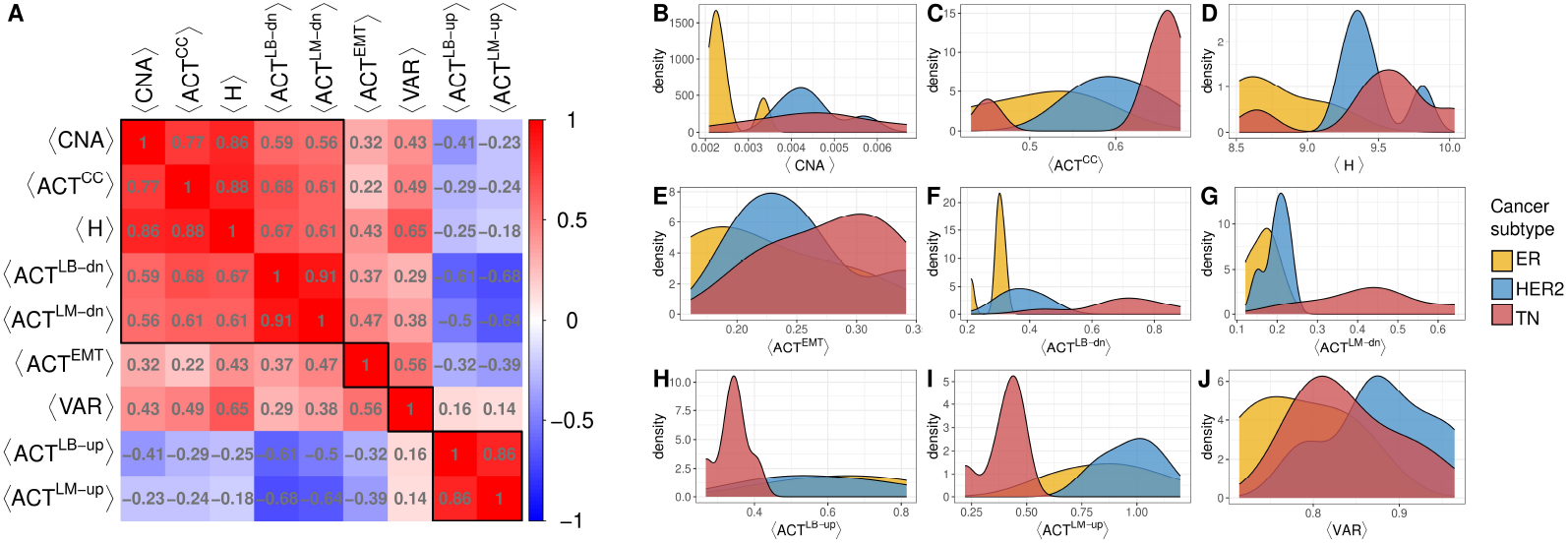
(A) Correlation matrix (Pearson correlation coefficient) between the nine sample parameters computed: ⟨*CNA*⟩, ⟨*ACT*^*CC*^⟩, ⟨*H*⟩, ⟨*ACT*^*LB*-*dn*^⟩, ⟨*ACT*^*LM*-*dn*^⟩, ⟨*ACT*^*EMT*^⟩, ⟨*V AR*⟩, ⟨*ACT*^*LB*-*up*^⟩, and ⟨*ACT*^*LM*-*up*^⟩. Red indicates a positive correlation, blue indicates a negative correlation, and white represents no correlation between the scores. The black squares identify the clusters obtained using hierarchical clustering. (B-J) Distribution estimation of sample scores, color-coded according to cancer subtypes: ER+, HER2+, and TN.

These findings suggest that samples with higher CNA burden tend to display increased cycling activity and entropy. Additionally, these samples manifest a basal and mesenchymal-like phenotype, potentially indicative of the cancer cells’ capability for EMT in more aggressive tumors. Intratumor transcriptomic heterogeneity ⟨*V AR*⟩, was also positively correlated with these previous scores; however, surprisingly, it also showed a positive correlation with ⟨*ACT*^*LM*-*dn*^⟩ and ⟨*ACT*^*LB*-*dn*^⟩, albeit with lower values.

Analysis of the nine sample score distributions across breast cancer subtypes revealed distinct patterns. ⟨*CNA*⟩ exhibited greater heterogeneity between the samples of TN compared to ER+ and HER2+ samples. Additionally, ⟨*CNA*⟩ levels were elevated in HER2+ compared to ER+, as visualized in figure 5B. Furthermore, scores ⟨*ACT*^*CC*^⟩, ⟨*H*⟩, ⟨*ACT*^*EMT*^⟩, ⟨*ACT*^*LB*-*dn*^⟩ and ⟨*ACT*^*LM*-*dn*^⟩ demonstrated the highest levels in TN samples. HER2+ samples displayed intermediate levels, followed by ER+ samples, as visualized in figures 5C-G. This trend aligns with the malignancy levels across cancer subtypes, where higher scores correspond to a more unfavorable prognosis. While there are multiple classifications of breast cancer subtypes, there is a consensus among the analyzed subtypes in this study that the order from better to worse prognosis would be ER+, HER2+, and TN [72–75]. In figures 5H and I, the opposite distribution order is observed. TN corresponds to a basal/mesenchymal phenotype leading to lower activity related to luminal cancer types, while ER+ and HER2+ samples exhibit higher activity of these PPINs. However, it is important to note that these kernel density estimations are based on a limited number of samples, leading to estimated distributions with peaks that may not accurately represent the true underlying distributions. These peaks correspond to samples that deviate significantly from the general behavior, as we will discuss later.

In terms of sample variability, ER+ presented the most left-skewed distribution, HER2+ the most right-skewed distribution and TN distribution fell between them (figure 5J). An explanation for this finding could be that HER2+ tumors present both luminal and basal features [73], and therefore exhibit heterogeneous transcriptomic patterns. This observation suggests that even though variability correlates positively with other scores (albeit to a lesser extent), this score does not necessarily indicate more aggressive tumors. In various cancer types, such as breast cancer, a distinct subset of cells known as cancer stem cells (CSCs) has been identified. These cells constitute only a small fraction [65,76], approximately 0.1 − 1%, of the total tumor cell population [77] and have been shown to contribute to poor patient prognosis. However, regarding the transcriptomic variability score, these cells may not exert a significant impact due to their low numerical abundance.

For a detailed inspection, in figure 6 we presented the scatter plots of selected sample parameters to observe their relationships, distinguishing them by cancer subtype and labeling the samples. The positive correlation between the scores reported in the correlation matrix (figure 5A) can be verified by these scatter plots. Moreover, certain general trends can be observed. ER+ samples tend to cluster at the lower regions of the scatter plots, indicating lower scores. In contrast, HER2+ and TN samples are more difficult to discriminate between each other, yet there is a tendency for TN samples to cluster at higher score levels compared to HER2+ samples.

**Figure 6.**
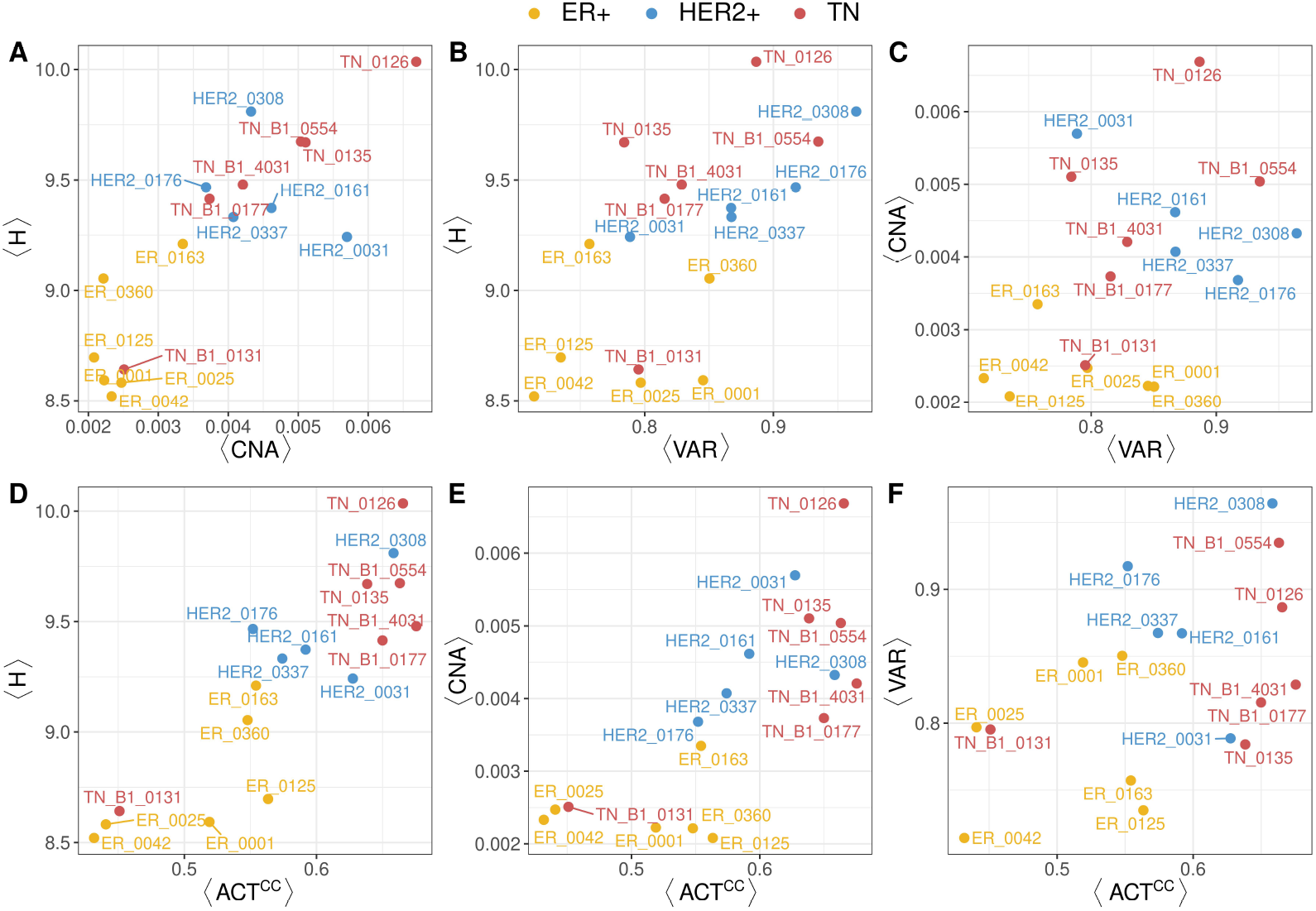
(A-F) Scatter plots illustrating several sample parameters that support the positive correlation. Each data point represents a sample, color-coded based on the cancer subtype, with labels included alongside the data points.

Examining samples individually reveals certain exceptions. For instance, among the TN samples, TN-B1-0131 displays low scores across all parameters except for ⟨*V AR*⟩, resembling profiles found in ER+ samples. Further scrutiny of the metadata reveals that the TN-B1-0131 sample corresponds to an 84-year-old patient (see Table 2.1), standing out as 20 years older than the subsequent TN sample when sorted by age. Additionally, it exhibits an age gap of more than 30 years compared to the average age of the other TN samples, which is 42. A similar observation in the opposite direction can be made for HER2-0308. This particular sample shows notably higher scores compared to the others of the same cancer subtype, except for ⟨*CNA*⟩. The donor for this sample is 32 years old (Table 2.1), younger by over 30 years than the mean HER2+ sample age (63). The exceptional score values observed may be linked to age-dependent tumor aggressiveness. Numerous studies have reported a more aggressive tumor biology, increased risk of recurrence, treatment failure rates, and mortality for younger patients [78–81]. The residual expression matrices displayed in figure 2 provide complementary information regarding *CNA*. In contrast to other TN samples, TN-B1-0131 exhibited a low ⟨*CNA*⟩ value. This is reflected in its residual expression matrix shown in figure 2C, which displays a pattern with fewer chromosomal gains and losses compared to the other TN samples. While HER2-0308 displayed intermediate levels of ⟨*CNA*⟩ within the HER2+ group (figure 2B), its residual expression matrix deviated from other HER2+ samples. Notably, it exhibited marked amplifications in chr6 and chr17, alongside deletions in chr14.

## 4. Conclusions

This study presents a quantitative approach to assess key features pertinent to cancer, specifically focusing on the three most prevalent subtypes of breast cancer: ER, HER2+ and TN. Using scRNA-seq data obtained from human breast cancer samples, we undertake a comprehensive examination of various cellular characteristics such as CNAs, entropy, and PPIN activity linked to specific biological processes (EMT, cell cycle, luminal, mesenchymal and basal breast cell lines). This approach contrasts with previous studies which concentrated on individual aspects or relied on qualitative analyses based on gene markers. At sample level, we computed the average of these measures. Additionally, we introduced a score that quantifies the sample intratumor transcriptomic heterogeneity.

The novelty of this study lies in the quantitative assessment of these features, a comprehensive approach previously unexplored in the cancer single-cell transcriptomics field and in particular in breast cancer.

Our investigation at the single-cell level has unveiled intriguing patterns. The PPIN activity associated with the cell cycle and entropy demonstrates varying degrees of activity across breast cancer subtypes: ER+, HER2+, and TN, in ascending order. Notably, clusters of cells displaying heightened mitotic activity are observed across all subtypes, with TN samples exhibiting a higher proportion of mitotic cells, indicating particularly pronounced activity. Our study revealed distinct CNA distribution patterns between ER+ and HER2+/TN tumors. HER2+/TN tumors exhibited significantly higher CNA values and greater dispersion compared to ER+ tumors. Furthermore, the activity profiles linked to basal and luminal cell lines employed in our study differentiate between basal and luminal tumors. We did not observe cell groups with distinctive elevated *ACT*^*EMT*^ in any subtype. This could be due to the reported scarcity of cells undergoing EMT, which may be masked within the large pool of cells analyzed [82]. Additionally, low expression levels of EMT-related genes, such as ZEB1, ZEB2 and SNAIL, might be sufficient to trigger EMT even without a substantial increase in *ACT*^*EMT*^. Another factor could be the absence of metastasis in the samples studied, leading to minimal EMT activity.

In our analysis of the sample mean scores, we identified a positive correlation among ⟨*CNA*⟩, ⟨*ACT*^*CC*^⟩, ⟨*H*⟩, ⟨*ACT*^*LB-dn*^⟩, and ⟨*ACT*^*LM-dn*^⟩, indicating that samples exhibiting basal characteristics present higher levels of these parameters. Moreover, these parameters show increasing levels in ER+, HER2+, and TN tumors (in ascending order), consistent with the known aggressiveness of these subtypes, except for ⟨*CNA*⟩. ⟨*CNA*⟩ was higher in HER2+ and TN compared to ER+ samples, but did not show a significant difference between HER2+ and TN tumors. While ⟨*ACT*^*EMT*^⟩ correlated positively with the previous scores, its overall correlation coefficient values were lower than among them. Additionally, the distributions of ⟨*ACT*^*EMT*^⟩ in ER+, HER2+, and TN samples exhibited a trend of increasing bias towards higher values, in that order.

An interesting exception emerged regarding ⟨*V AR*⟩, a parameter quantifying transcriptomic heterogeneity within each sample, where we observed higher values for HER2+, followed by TN, and lastly ER+ tumors. This observation can be explained by the fact that HER2+ tumors often possess both luminal and basal properties, resulting in a more diverse range of transcriptomic profiles.

Our quantitative measures uncovered distinct behavior in samples HER2-0308 and TN-B1-0131 compared to the others within their respective subtypes. This difference could be attributable to a patient age disparity relative to the age range of the rest of the samples for each subtype.

## Ethics

This work did not require ethical approval from a human subject or animal welfare committee.

## Data Accessibility

Data used in this work are available in the GEO database under the accession number GSE161529 and on [figshare](https://figshare.com/articles/dataset/Data_R_code_and_output_Seurat_Objects_for_single_cell_RNA-seq_analysis_of_human_breast_tissues/17058077). All relevant code has been deposited in in the following GitHub repositories: https://github.com/danielasenraoka/R-code-workflow-Unraveling-tumor-heterogeneity and https://github.com/danielasenraoka/PyOrigins.

## Authors’ Contributions

D.S.: Conceptualization; Data curation; Formal analysis; Investigation; Methodology; Software; Validation; Visualization; Writing - original draft. N.G.: Conceptualization; Funding acquisition; Resources; Supervision; Writing - review & editing. L.D.: Conceptualization; Formal analysis; Funding acquisition; Project administration; Resources; Software; Supervision; Writing - review & editing.

## Acknowledgements

We thank Martín Tomé for his technical support and for providing valuable insights. His expertise and assistance were extremely helpful in the execution of this project.

