## Supplementary Figures for "Unraveling tumor heterogeneity: Quantitative insights from scRNA-seq analysis in breast cancer subtypes"

### Supplementary Materials

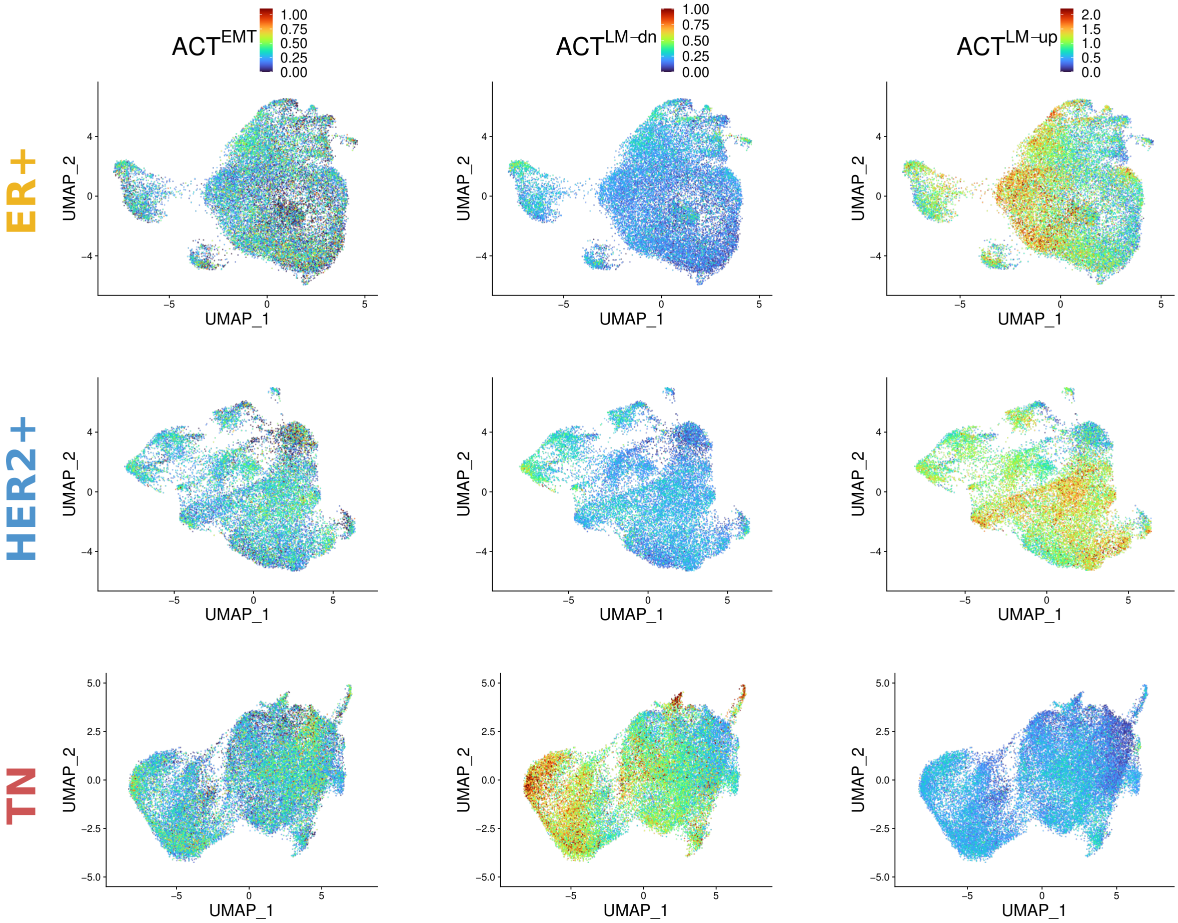


**Figure S1:** UMAP representation of integrated samples separated by cancer subtype: ER+, HER2+, and TN. Cells are color-coded according to the scores $ACT^{EMT}$, $ACT^{LM-dn}$, and $ACT^{LM\text{-}up}$.
